# Oxidative Stress is Intrinsic to Staphylococcal Adaptation to Fatty Acid Synthesis Antibiotics

**DOI:** 10.1101/2022.09.09.506511

**Authors:** Paprapach Wongdontree, Aaron Millan-Oropeza, Jennifer Upfold, Jean-Pierre Lavergne, David Halpern, Clara Lambert, Adeline Page, Gérald Kénanian, Christophe Grangeasse, Céline Henry, Agnès Fouet, Karine Gloux, Jamila Anba-Mondoloni, Alexandra Gruss

## Abstract

Antibiotics inhibiting the fatty acid synthesis (FASII) pathway of the major pathogen *Staphylococcus aureus* reach their enzyme targets, but bacteria continue growth by using environmental fatty acids (eFAs) to produce phospholipids. We assessed how extreme changes in membrane phospholipids provoked by FASII-antibiotics affect global *S. aureus* physiology. Anti-FASII provoked massive lasting expression changes without genomic rearrangements. Several regulators, rather than one master switch, contributed to the timing of anti-FASII adaptation. Numerous virulence and adhesion factors showed decreased levels and/or activity. Conversely, stress response protein levels increased, and correlated with greater tolerance to peroxides. Notably, peroxide priming stimulated eFA incorporation efficiency and facilitated adaptation to FASII inhibition. These findings establish a link between oxidative stress and FA incorporation. Consistent with major shift in protein expression, anti-FASII-adapted *S. aureus* killed an insect host more slowly but continued multiplying. Thus, while anti-FASII-adapted populations are less equipped to damage the host, they may be better fit for long term survival, and could constitute a reservoir for re-infection.

## Introduction

*Staphylococcus aureus* is a Gram-positive opportunistic bacterium that remains a major cause of disease and mortality in humans and animals. The unsolved crisis of non-treatable infections due to multidrug-resistance notably methicillin-resistant *S. aureus* (MRSA) underlines the need for alternative treatments, especially in compromised patients (Lowy, 1998, Rasigade and Vandenesch, 2014, Lakhundi and Zhang, 2018, Cusumano et al., 2020).

Enzymes of the fatty acid (FA) synthesis pathway (FASII) are longtime candidate targets for drug development against *S. aureus* infections (Wang et al., 2006, Wright and Reynolds, 2007). FabI, enoyl-ACP reductase, was a preferred target as a narrow spectrum inhibitor that would not disrupt the gut microbiome during treatment (Karlowsky et al., 2009). However, while anti-FASII drugs targeting FabI effectively reached their targets in *S. aureus*, bacterial growth continued in *in vitro* or mouse bacteremia models; this is because bacteria incorporate environmental FAs (eFAs), which are abundant in host biotopes and are used directly to produce bacterial membrane phospholipids; the process is termed FASII bypass (Brinster et al., 2009, Gloux et al., 2017, Kenanian et al., 2019, Morvan et al., 2016) (**Fig. S1** schematizes FASII and FASII bypass pathways). Exposure to host lipids during septicemic infection thus favors *S. aureus* adaptation to anti-FASII treatment (Kenanian et al., 2019). Despite these limitations of FASII antibiotics in treating streptococcal, enterococcal, and staphylococcal infections, new anti-FASII drugs are in development (Pishchany et al., 2018, Yao and Rock, 2016, Fage et al., 2020, Parker et al., 2022). Characterizing the anti-FASII-adapted *S. aureus* populations that might persist in the host after treatment is thus crucial for future use.

We report here that anti-FASII treatment results in the emergence of a novel *S. aureus* fitness state. Anti-FASII-adapted bacteria produce less virulence factors. In contrast, stress responses are activated, and adapted bacteria show greater peroxide tolerance. Pre-exposure to peroxides accelerates anti-FASII adaptation by enhancing fatty acid incorporation, thus identifying a new strategy by which peroxide priming facilitates antibiotic survival. These findings elucidate the state of anti-FASII-adapted bacteria that persist in the host after anti-FASII treatment.

## Results

### Anti-FASII treatment leads to long-term *S. aureus* adaptation without detectable chromosomal rearrangements

Our previous studies ruled out a role for point mutations or indels in *S. aureus* adaptation to anti-FASII in serum-containing medium; studies were performed using either of two potent FabI inhibitors, triclosan (McMurry et al., 1998) or AFN-1252, a pipeline antimicrobial with high FabI specificity (Karlowsky et al., 2009), giving equivalent results (Kenanian et al., 2019). As large chromosomal rearrangements are not detected by mi-Seq (Kenanian et al., 2019), we resorted to nanopore sequencing of four anti-FASII-adapted cultures. Tests were performed using AFN-1252 and two non-treated cultures. No rearrangements specific to anti-FASII adaptation were detected, showing that anti-FASII-adaptation occurs in the absence of chromosomal rearrangements or mutations. Once adapted to anti-FASII, growth was robust in SerFA liquid medium (BHI supplemented with serum and fatty acids); however, bacteria grew poorly in FA-free medium (**Fig. S2**). These results indicated that phenotypic alterations were responsible for anti-FASII adaptation.

### Anti-FASII adaptation involves massive protein reprogramming

A proteomics approach was used to elucidate changes occurring during anti-FASII adaptation. FASII inhibition is overcome by eFAs provided in the presence of serum. Serum protects bacteria from toxic free FAs, and promotes FASII bypass (Kenanian et al., 2019). A complete kinetics study on *S. aureus* USA300 used the anti-FASII triclosan, both without and with added serum, at 2, 4, 6, 8, and 10 h post-anti-FASII addition (Fig. 1A, B, C respectively show strategy, growth kinetics, and proteome results; **Table S1, tabs 1 and 2,** give complete results according to cluster analyses). As expected, protein abundance was mainly lower in triclosan medium (FA-Tric) without serum, where adaptation occurs by mutation emergence (Morvan et al., 2016). In contrast, massive protein level changes in adaptation (SerFA-Tric) medium suggest metabolic activity in latency phase prior to full outgrowth, with transient or lasting expression differences compared to non-treated bacteria (compare SerFA and SerFA-Tric, Fig. 1C).

**Fig. 1.**
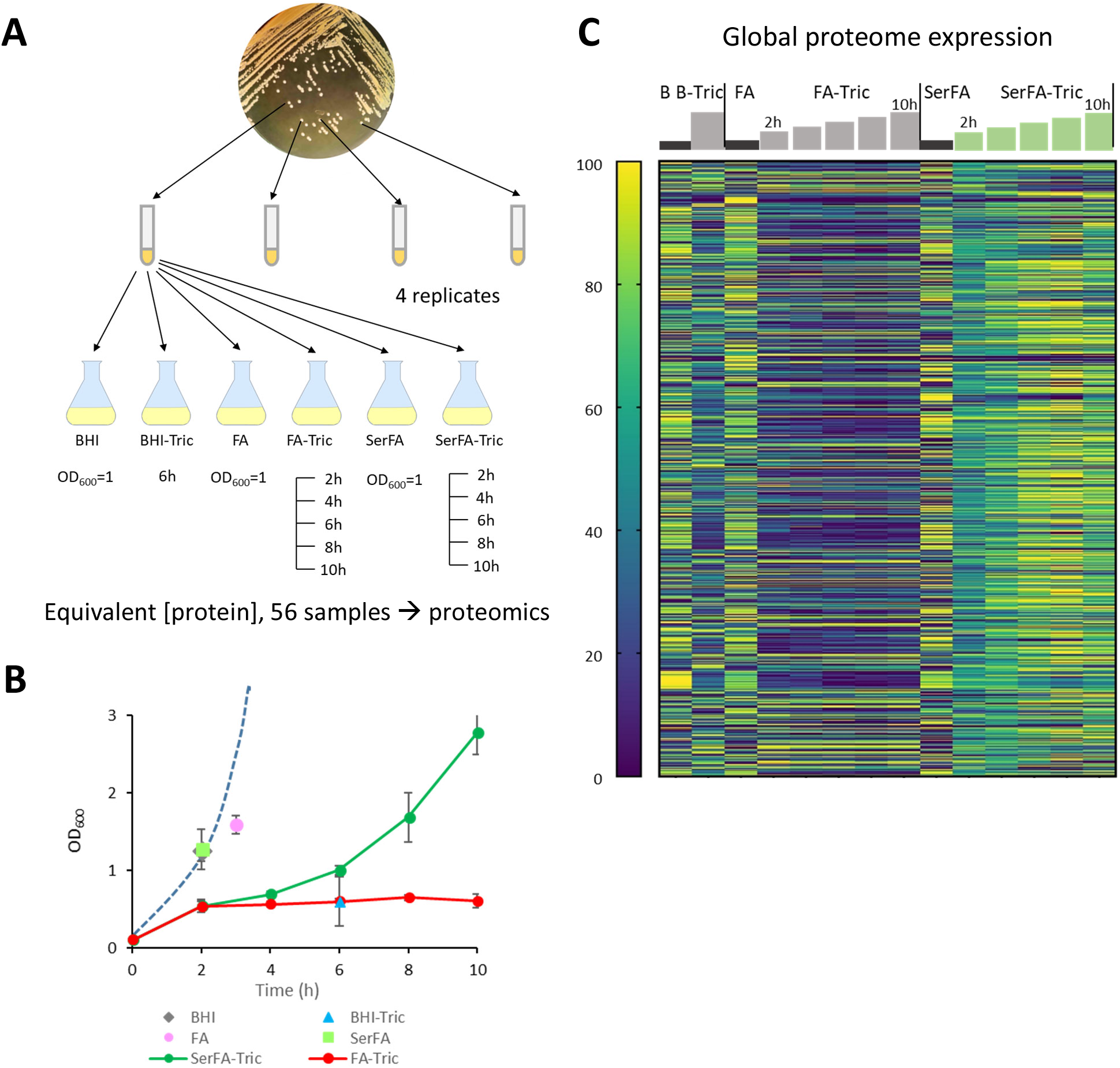
Proteomic kinetics and global expression changes related to *S. aureus* USA300 strain adaptation to anti-FASII. A. Schematics of sample preparation. Samples (4 biological replicates) were grown and prepared in conditions and harvesting times (or OD_600_ for controls) as indicated below flasks. **B. Sample growth.** Means and standard deviations are indicated. Points correspond to sampling times. See **Table S1** for details. Dashed blue line represents typical growth in SerFA medium determined separately. *S. aureus* adapts to anti-FASII within 10h on SerFA-Tric, but not on FA-Tric, after a latency period (double arrow). **C. Heat map of proteins showing altered levels in at least 1 condition relative to other samples.** All sample conditions are shown. Sampling times (h) above steps correspond to 2, 4, 6, 8, and 10 h for FA-Tric and SerFA-Tric. The heat map shows global protein changes, and is determined relative to weighted value for each protein, as on scale at left (navy, down-represented; yellow, up-represented).

We first exploited the data to screen for proteins whose levels selectively increased in the presence of specific signals related to FAs and/or anti-FASII (**Fig. S3**). Highest expression of several Type 7 secretion system proteins (comprised between SAUSA300_0278 and SAUSA300_0289) occurred in FA conditions, as reported (Lopez et al., 2017). The FarE fatty acid efflux pump was also up-regulated, presumably to avoid accumulation and toxicity of free FAs (Alnaseri et al., 2015); as reported, induction was alleviated in serum, which lowers toxicity (Kenanian et al., 2019). Overall, serum markedly affects *S. aureus* responses to anti-FASII, thus highlighting its crucial role in adaptation.

Proteins related to FASII and phospholipid metabolism showed striking differences according to conditions (**Fig. S4**). Levels of most FASII- and phospholipid-related enzymes increased in SerFA-Tric as compared to those in all other conditions, including FA-Tric without serum. Notably, PlsC (1-acyl-sn-glycerol-3-phosphate acyltransferase), is required for phospholipid synthesis and anti-FASII adaptation. After an initial decrease, PlsC levels are then restored during adaptation outgrowth, in accordance with results using a reporter fusion (Pathania et al., 2021). This result highlights the importance of serum in enabling FASII bypass.

We focus below on the changes occurring in serum-supplemented medium containing anti-FASII (SerFA), in which FASII bypass occurs without genetic modification. Three categories of proteins, regulators, stress response proteins, and virulence factors, are analyzed.

### Anti-FASII adaptation triggers shifts in regulatory protein expression and phosphorylation

Regulatory proteins are likely implicated in the massive changes caused by anti-FASII treatment. Proteomics revealed 22 hypothetical or confirmed proteins implicated in regulation whose expression decreased (9) or increased (13) during anti-FASII adaptation; they affect stress response, virulence, nutritional immunity and metabolism, or combinations of these factors (Fig. 2A). As phosphorylation is a frequent post-translational modification that alters functions of numerous regulatory proteins (Garcia-Garcia et al., 2016, Mijakovic et al., 2016, Derouiche et al., 2013), we performed a phosphoproteomics analysis, focusing here on regulatory proteins. Five regulators showed altered phosphorylation patterns during transition or upon adaptation to anti-FASII (Fig. 2B and **Fig. S5**; **Table S2** for full results). Altogether, we selected CcpE, CshA, HrcA, Rot, and XdrA as showing high expression and/or phosphorylation changes at or just prior to adaptation outgrowth (8-10 h) (Fig. 2A and B) for further study.

**Fig. 2.**
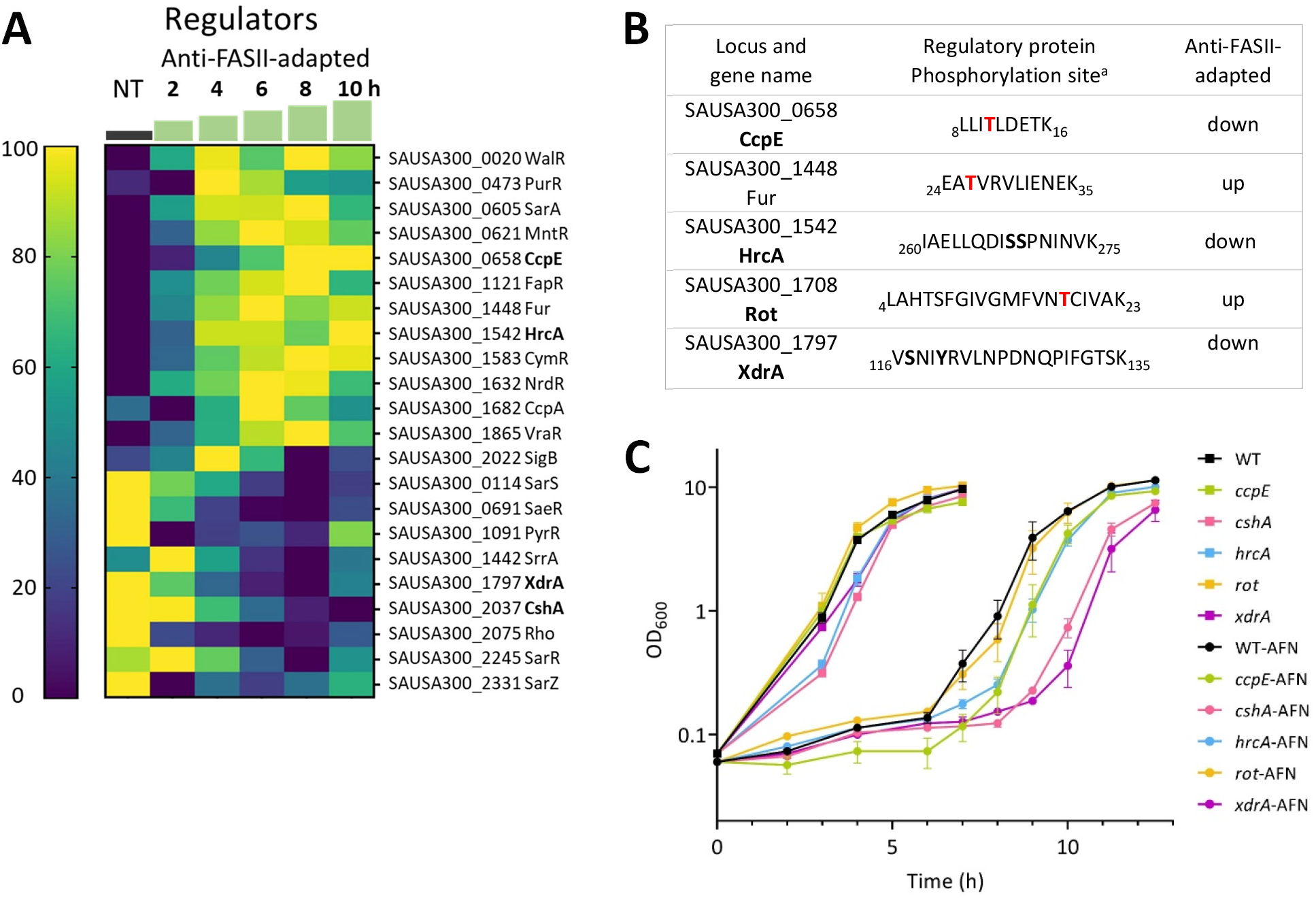
Anti-FASII affects pools and phosphorylation status of regulatory proteins. A. Expression of known or putative regulators that differ in at least one time point during anti-FASII adaptation (SerFA-Tric) compared to non-treated (NT) SerFA sample. Changes in putative regulatory protein levels are shown in anti-FASII adaptation SerFA medium (see **Table S1** for all test conditions). Gene loci and names are at right. Times (h) of sampling are indicated above green steps. Heat map (scale at left) is determined relative to weighted value for each protein (navy, down-represented; yellow, up-represented). **B**. **Phosphorylated regulatory proteins with altered expression during anti-FASII adaptation.** Peptides showing differential phosphorylation at 10 h post anti-FASII adaptation are indicated (see **Table S2** for complete information). ^a^ Peptide positions in respective protein sequences are indicated. Phosphorylated amino acids are in bold red or in bold black when there is an ambiguity. **C. Effects of regulator gene inactivation on anti-FASII adaptation.** Kinetics of anti-FASII-adaptation of USA300 (WT) and insertional mutants, based on regulators with altered protein and/or phosphorylation levels (**A** and/or **B**; selected loci in bold), were compared to that of the parental strain. Cultures were grown without (squares) and with anti-FASII AFN-1252 (circles). Growth curves and standard deviations are based on biological triplicates.

To determine how these regulators affected anti-FASII adaptation, we compared growth kinetics of USA300 and the corresponding mutant strains (Fey et al., 2013) (Fig. 2C). The absence of *cshA* led to ∼2 h delays in anti-FASII adaptation time. CshA controls the supply of acetyl-CoA; its interruption results in phospholipids enriched in saturated FAs (Khemici et al., 2020). We propose that this membrane alteration might interfere with eFA incorporation.

The *xdrA* mutant also showed a 2 h delay in adaptation. Both the amounts and phosphorylation of XdrA decreased during anti-FASII treatment. The putative fatty acid degradation gene *fadX* is reportedly upregulated when XdrA pools are lower (McCallum et al., 2010), which might transiently compete for eFAs and limit their availability for FASII bypass. Alternatively, XdrA DNA binding activity reportedly overlaps that of the CodY regulator (Lei and Lee, 2018). We recently showed that a *codY* mutant is delayed in adaptation, which we proposed relates to a positive regulatory effect of CodY on Acc expression (Pathania et al., 2021). The delay in anti-FASII adaptation of the *xdrA* mutant might relate to a coordinate role between XdrA and CodY. *hrcA* and *ccpE* mutants showed minor changes in anti-FASII adaptation kinetics, while *rot* had no detectable change (Fig. 2C).

Thus, while regulatory mutants individually affect anti-FASII-adaptation efficacy, no single tested regulator acted as a master controller. In *Escherichia coli*, stress response regulators respond to discrete extracytoplasmic signals, each of which independently contributes to stress adaptation (Bury-Mone et al., 2009). Similarly, differentially expressed regulators may each respond to different stimuli consequent to membrane perturbation by anti-FASII, which cooperatively promote adaptation.

### Reduced levels and activities of virulence factors in anti-FASII-adapted *S. aureus*

Levels of 10 out of 16 detected virulence-related proteins decreased transiently or durably during anti-FASII adaptation compared to non-treated *S. aureus* cultures (Fig. 3A). Levels of several adhesins were markedly decreased, including ClfB, which is required for virulence (Dyzenhaus et al., 2022); iron receptor protein and toxin levels were also overall reduced. In contrast, two peptidoglycan hydrolases, Atl and IsaA, showed increased levels. Hydrolases are implicated in housekeeping, and also in virulence (Stapleton et al., 2007); their increased expression upon adaptation outgrowth might resolve cell wall thickening in the latency phase (PW, AG, JAM, in preparation). We asked whether decreased amounts of adhesion factors SdrD, SdrE, EbpS, and ClfB in anti-FASII-adapted *S. aureus* might impact adhesion to host cells. Anti-FASII-adapted *S. aureus* (using AFN-1252) adhered poorly to human THP-1 macrophages compared to non-adapted bacteria (20% *versus* 44%, p < 0.01; Fig. 3B). We also evaluated major secreted virulence factors not detected by proteomics: lipase, nuclease, protease, and hemolysin in non-treated *versus* anti-FASII-adapted *S. aureus* cultures (Fig. 3C). Activities of these exoproteins were visibly lower in anti-FASII adapted cultures obtained using triclosan or AFN-1252. Altogether these results indicated that anti-FASII-adapted *S. aureus* populations may be less fit for virulence.

**Fig. 3.**
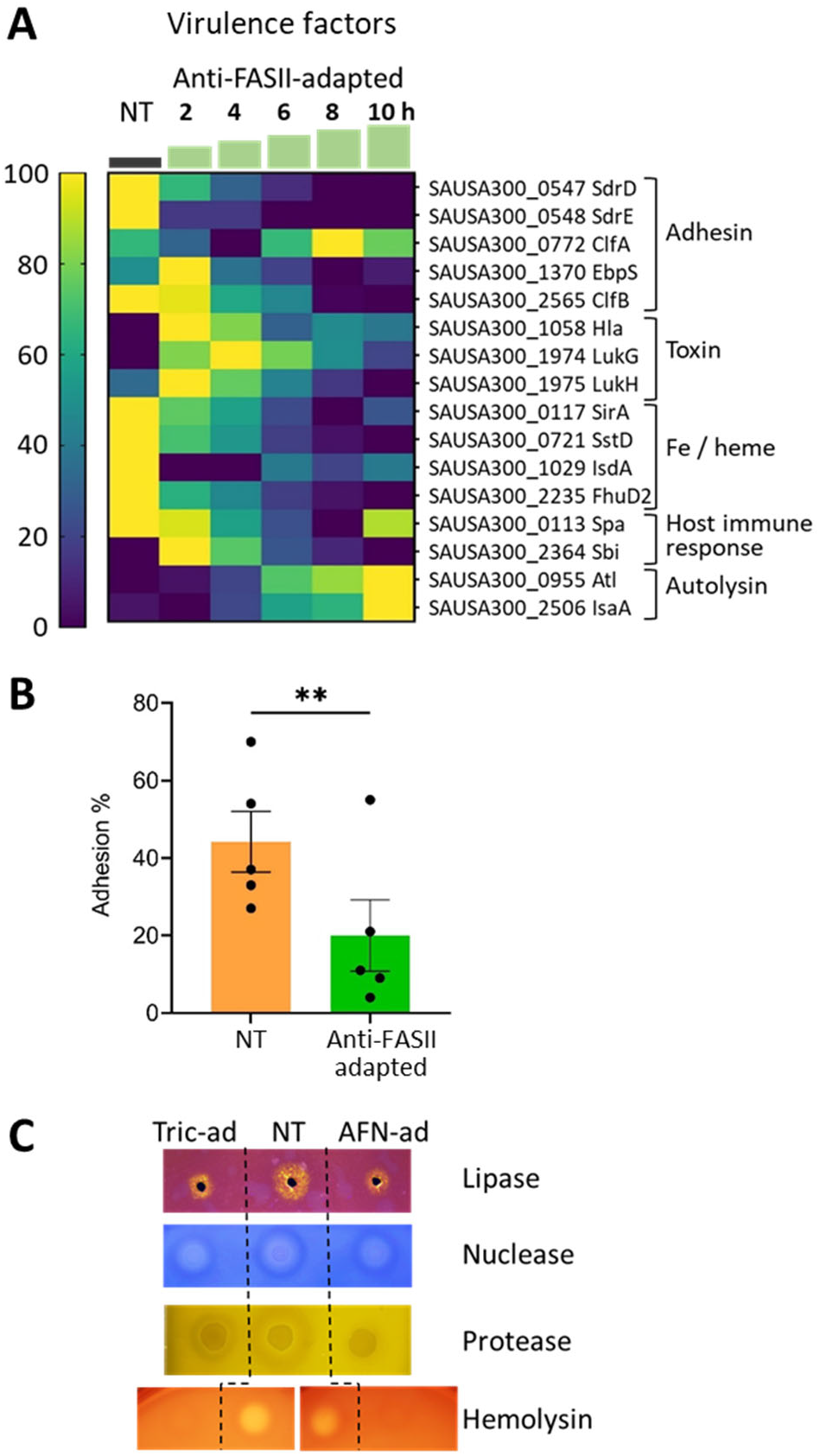
Expression changes during anti-FASII treatment. A. Heat maps of known or putative virulence factors. Samples shown are in anti-FASII adaptation medium (SerFA; see **Table S1** for results in all test conditions). Kinetics (in h) of sampling is indicated above green steps. Gene loci, protein names, and functional categories are at right. Correspondence between color and expression in heat map (scale at left) is determined relative to weighted value for each protein (navy, down-represented; yellow, up-represented). **B. Differential NT and anti-FASII-adapted *S. aureus* adhesion to macrophage.** Non-treated (NT) or AFN-1252-treated *S. aureus* USA300 (3×10^5^) were added to of THP-1 macrophage monolayers (3×10^5^ cells per well), and incubated for 1 h at 4° C. Colony forming units (CFU) of bacteria that adhered to macrophage were determined on five independent bacterial samples (P= 0.003; see Materials and Methods). **C. Secreted virulence factor activities.** Non-treated (NT) and anti-FASII-adapted cultures treated with triclosan (Tric-ad) or AFN-1252 (AFN-ad) were grown overnight, and reached similar OD_600_ values (= 13, 9 and 9 respectively for NT, Tric-ad, and AFN-ad). Cultures (for protease detection) and culture supernatants (lipase, nuclease, and hemolysin detection) were prepared (see Materials and Methods) and spotted on appropriate detection medium. Representative results of 3 biologically independent replicates are shown.

### Anti-FASII-adapted *S. aureus* produce increased levels of stress response proteins and show increased resistance to an oxidative stress

Levels of 26 out of 28 assessed stress response proteins showed transient or lasting increases in anti-FASII-adapted *S. aureus* compared to the non-treated control. Among them, 18 were pronounced in during adaptation outgrowth. Proteins are implicated in pH, oxidative, osmotic, and unfolded protein responses (Fig. 4A).

**Fig. 4.**
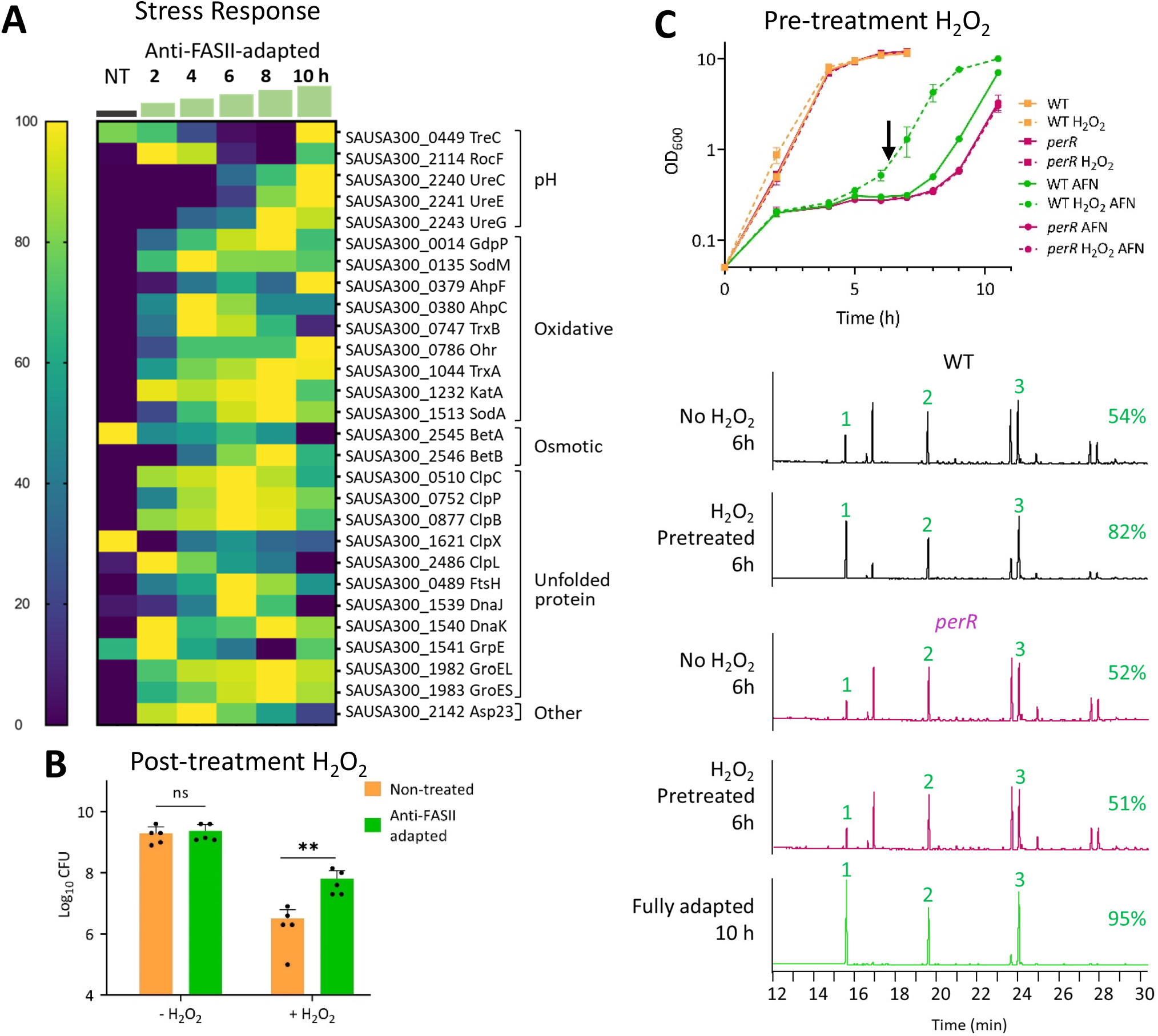
Anti-FASII adaptation confers oxidative stress resistance, and is accelerated by prior exposure to peroxide stress. A. Stress response heatmap. Results are shown for anti-FASII adaptation medium (SerFA; see **Table S1** for all test conditions). Sampling times (h) are indicated above green steps. Gene names and functional categories are at right. Heat map (scale at left) is determined relative to weighted value for each protein (navy, down-represented; yellow, up-represented). B. *S. aureus* anti-FASII adaptation confers increased H_2_O_2_ resistance. USA300 non-treated (NT, orange bar) and AFN-1252-adapted overnight cultures (green bar) were challenged with 0.5 mM H_2_O_2_ for 5 h, and CFUs were determined. **, p< 0.01. C. Priming *S. aureus* with H_2_O_2_ accelerates anti-FASII adaptation and requires PerR. Upper: USA300 (WT) and *perR* (SAUSA300_1842) mutant SerFA cultures were grown overnight without or with 0.5 mM H_2_O_2_. Cultures were diluted (OD_600_ = 0.1) in SerFA without or with 0.5 µg/ml AFN-1252, and growth was monitored. Results are shown for biological triplicates. H_2_O_2_ – primed samples, without or with AFN addition at T0 are as indicated. **Lower:** FA profiles of indicated strains harvested after 6 h anti-FASII treatment (arrow in ‘C’). In green, FA profile of fully adapted H_2_O_2_-pretreated cultures at 10 h, shown here for *perR* and non-distinguishable from WT. 1, 2, and 3, eFAs present in SerFA medium (respectively C14, C16, and C18:1). At left of each profile, proportion of incorporated eFAs (%) are the average of two independent measurements (<3% difference between replicates).

Host production of reactive oxidative species (ROS) is a documented response to infection as a means to eliminate the pathogen (Liu et al., 2005). We asked whether increased levels of stress-related factors in anti-FASII-adapted *S. aureus* affected bacterial tolerance to ROS. This was tested by challenging non-treated and anti-FASII-adapted cultures with 0.5 mM H_2_O_2_ for 5 h, followed by survival assessments by colony forming unit (CFU) determinations. Anti-FASII-adapted *S. aureus* CFUs were ∼20-fold higher than for non-treated bacteria (p <0.01; Fig. 4B). Greater peroxide tolerance might thus confer a survival advantage to the anti-FASII-adapted, oxidative stress-resistant populations.

### Priming with peroxides prior to anti-FASII challenge stimulates adaptation by enhancing eFA incorporation, and requires PerR

Stress response induction by anti-FASII led us to ask the contrary, whether stress priming would stimulate anti-FASII bypass. A ROS priming effect reportedly increased resistance to various antibiotics in *E. coli*, including oxolinic acid, ampicillin, kanamycin (Mosel et al., 2013), and fluoroquinolones (Gerstel et al., 2020). The effects of ROS priming on anti-FASII adaptation in *S. aureus* were tested by subjecting cultures to overnight priming with sub-lethal concentrations of H_2_O_2_ (0.5 mM) or phenazine-methosulfate (PMS, a redox cycling compound; 20-50 µM (Gerstel et al., 2020)). Cultures were diluted, and kinetics of anti-FASII (AFN-1252) adaptation was followed. H_2_O_2_ or PMS priming shortened the time of *S. aureus* adaptation to FASII antibiotics by 1.5 – 1.7 h compared to non-primed WT cultures (Fig. 4C **upper**, Table 1). A *katA* mutant lacks catalase and would accumulate H_2_O_2_; anti-FASII adaptation in *katA* was ∼1 h shorter than in the WT parental strain. Priming *katA* with H_2_O_2_ as above further shortened the adaptation time by ∼2 h as compared to WT without peroxide (Table 1). Conversely, treating the WT strain with reducing agents (Na citrate 10 mM, or vitamin C 5.7 mM) retarded adaptation by 3.5 and 1.7 h respectively (Table 1). *S. aureus* oxidation state is thus implicated in anti-FASII adaptation.

**Table 1.** Pre-treatments with ROS-generating *versus* reducing agents respectively accelerate and retard adaptation to anti-FASII.

| Strain<sup>a</sup> | Additive<sup>b</sup> | Time (h) to exit | $\Delta$ anti-FASII exit (h) compared to WT (SD)<sup>c</sup> | N |
| --- | --- | --- | --- | --- |
| WT | - | 8 | - | 8 |
| WT | H<sub>2</sub>O<sub>2</sub> | 6.3 | + 1.68 (±0.45) | 8 |
| WT | PMS | 6.5 | + 1.50 (±0.35) | 5 |
| *katA* | - | 7 | + 1.00 (±0.00) | 3 |
| *katA* | H<sub>2</sub>O<sub>2</sub> | 5.8 | + 2.17 (±0.58) | 3 |
| *perR* | - | 9 | - 1.00 (±0.00) | 3 |
| *perR* | H<sub>2</sub>O<sub>2</sub> | 9 | - 1.00 (±0.00) | 3 |
| *sufD* * | - | 7.3 | + 0.75 (±0.50) | 4 |
| *sufD* * | H<sub>2</sub>O<sub>2</sub> | 6.7 | + 1.25 (±0.29) | 4 |
| WT | Na citrate | 11.5 | - 3.50 (±0.71) | 2 |
| WT | Vit C | 9.8 | - 1.75 (±0.35) | 2 |

<sup>a</sup> Strains USA300 (WT) or mutant derivatives *katA* SAUSA300\_1232, *perR* SAUSA300\_1842, or *sufD*\* intergenic between *sufC* (SAUSA300\_0818) and *sufD* (SAUSA300\_0819), at position 898566 (Fey et al., 2013, Roberts et al., 2017), are used. <sup>b</sup> Oxidizing agents were added only to precultures: H<sub>2</sub>O<sub>2</sub>, 0.5 mM; phenazine-methosulfate (PMS), 20 (n=2), 30 (n=1), 40 (n=1), 50 (n=1) μg/ml, gave equivalent results. Reducing agents were added at the same time as anti-FASII, here AFN-1252: sodium citrate, 10 mM; vitamin C (VitC), 5.7 mM. <sup>c</sup> Bacterial outgrowth for USA300 in SerFA-AFN is observed at 8 h, and is used as reference. Differences ( $\Delta$ ) in time of outgrowth are relative to that of the WT. +, early exit from latency (h); -, delayed exit (h). SD, standard deviation; N, number of independent experiments.

ROS priming induces antibiotic efflux in *E. coli*, leading to antibiotic tolerance (Gerstel et al., 2020, Mosel et al., 2013, Wu et al., 2012). Induction of the *E. coli* Fe-S repair system Suf matures the SoxR regulator, which in turn activates the AcrAB efflux pump to eliminate fluoroquinolones (Gerstel et al., 2020). We asked whether peroxide similarly induces an efflux mechanism that would accelerate anti-FASII adaptation. If this were the case, FASII activity would be restored, and bacteria would be anti-FASII-sensitive. However, examination of FA profiles revealed the contrary: at 6 h post anti-FASII treatment (using AFN-1252), *i.e.*, during the latency phase, H_2_O_2_ priming resulted in more efficient eFA incorporation compared to that in non-primed cultures (respectively 82% *versus* 54% eFAs; Fig. 4C). Moreover, a transposon insertion inactivating the *suf* homolog in *S. aureus* (Roberts et al., 2017) did not abolish the H_2_O_2_ priming effect (Table 1). These results show that stimulated eFA incorporation and hence more rapid adaptation, rather than anti-FASII efflux, explains the priming effects of peroxide on anti-FASII adaptation.

PerR (SAUSA300_1842) is a conserved highly sensitive metal-dependent peroxide sensor and regulator protein (Ji et al., 2015). PerR was not detected in proteomics of anti-FASII adaptation (**Table S1**). However, as PerR senses H_2_O_2_, we examined its implication in H_2_O_2_-accelerated anti-FASII adaptation. Unlike the USA300 parent, *perR* mutant adaptation time and FA incorporation efficiency were unaffected by peroxide (proportions of eFA incorporation at 6 h were 52% and 51%, with and without H_2_O_2_ priming; Fig. 4C). Altogether these results indicate the existence of a unique H_2_O_2_ priming system that requires PerR. They reveal a novel link between peroxide and FA incorporation that facilitates adaptation to the FASII class of antibiotics independently of antibiotic efflux.

### Anti-FASII-adapted *S. aureus* are less virulent, yet persist in a *Galleria mellonella* infection model

Lower virulence factor production but greater stress resistance due to anti-FASII adaptation raises questions on the outcome of anti-FASII-adapted *S. aureus* infection. We compared anti-FASII-adapted *versus* non-treated *S. aureus* in an insect *G. mellonella* model, which allows the use of a large cohort. Moreover, larval hemocoel, like blood serum, is lipid-rich (Kazek et al., 2021). AFN-1252, as chosen for this study, was pharmacologically vetted for non-toxicity in the host (Hunt et al., 2016, Parsons et al., 2011); we showed previously that AFN-1252 treatment did not stop infection in a mouse model (Kenanian et al., 2019). Non-treated and AFN-1252-adapted *S. aureus* USA300 showed equivalent growth kinetics in SerFA *in vitro* (**Fig. S2**). Insects infected by 10^6^ anti-FASII-adapted bacteria were killed more slowly than those infected by equivalent CFUs of untreated bacteria, as monitored over 72 h post-infection (Fig. 5A). At T_48_, 95% of insects were killed by untreated *S. aureus*, compared to 30% killing with anti-FASII-adapted *S. aureus*. At T_72_, when all larvae infected by untreated *S. aureus* were dead, anti-FASII-adapted *S. aureus* killed over 60% of the insects. Thus, killing by anti-FASII-adapted *S. aureus* was delayed but not stopped.

**Fig 5.**
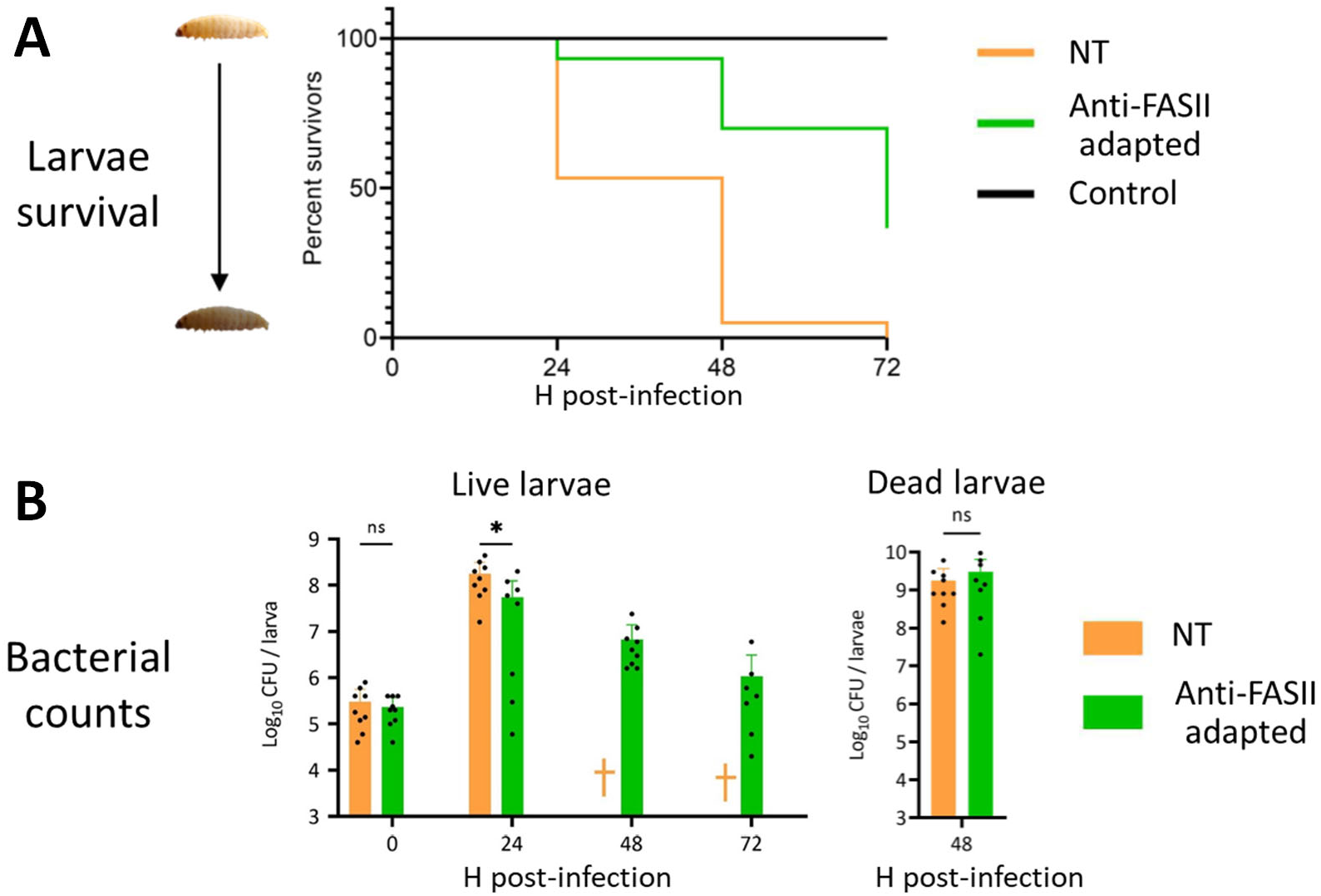
Comparison of untreated and anti-FASII-treated *S. aureus* in a *G. mellonella* infection model. Insects were injected with 10^6^ CFU *S. aureus* USA300 that were either non-treated (NT) or preadapted to anti-FASII (anti-FASII adapted) (see Materials and Methods). **A. Insect mortality in NT and anti-FASII-adapted *S. aureus*.** Data was analyzed using Kaplan-Meier with pooled values of biologically independent triplicates (60 insects per condition). Results excluding PBS controls (insects injected with equivalent volumes of PBS, black line) were analyzed by Mantel-Cox, and showed a significance value of p <0.0001. **B. Bacterial CFUs in surviving insects.** Insects were infected as in ‘**A**’. †, no surviving insects. **C. Bacterial CFUs in dead infected insects at 48 h.** ‘**B**’ and ‘**C**’ analyses were done using the non-parametric Mann Whitney test (Graphpad prism software). *, p = 0.02; ns, non-significant.

Bacterial CFUs determined from infected insects at 24 h were about 3-fold lower in those infected by anti-FASII-adapted compared to non-treated bacteria, and then decreased in surviving insects (Fig. 5B **left**). In contrast, CFUs in dead larvae at T_48_ were comparable for both groups, indicating that anti-FASII-adapted bacteria multiplied as well as non-treated bacteria in the insects they killed (Fig. 5B **right**).

To determine whether anti-FASII-adapted bacteria remained adapted during infection, CFU platings as above were done in parallel on SerFA and SerFA-AFN solid media. Only anti-FASII-adapted bacteria formed colonies on SerFA-AFN solid medium (**Table S3**). Among surviving insects infected with adapted bacteria, only one contained bacteria that returned to the non-adapted state at 72 h; this indicates that bacteria remained mainly adapted during insect infection. Altogether, these results show that anti-FASII-adapted *S. aureus* are less virulent, while populations still multiply in the insect host. Greater stress resistance capacity may help withstand host conditions and can explain bacterial persistence in the insect host.

## Discussion

eFA incorporation during FASII-antibiotic-induced bypass redesigns *S. aureus* membrane phospholipids, leading to massive shifts in protein expression, and a prolonged adaptation state without detectable genomic rearrangements or mutations (summarized in Fig. 6 model). These changes, which begin before bacterial outgrowth, alter the *S. aureus* fitness state towards less damage to the host, and more tolerance to host-generated toxicity. Peroxide priming accelerates adaptation and eFA incorporation, establishing a link between ROS and FASII bypass. The *in vitro* phenotypes deduced from proteomic and *in vitro* findings are consistent with the differences observed in insect infection kinetics between non-treated and anti-FASII-treated *S. aureus*.

**Fig 6.**
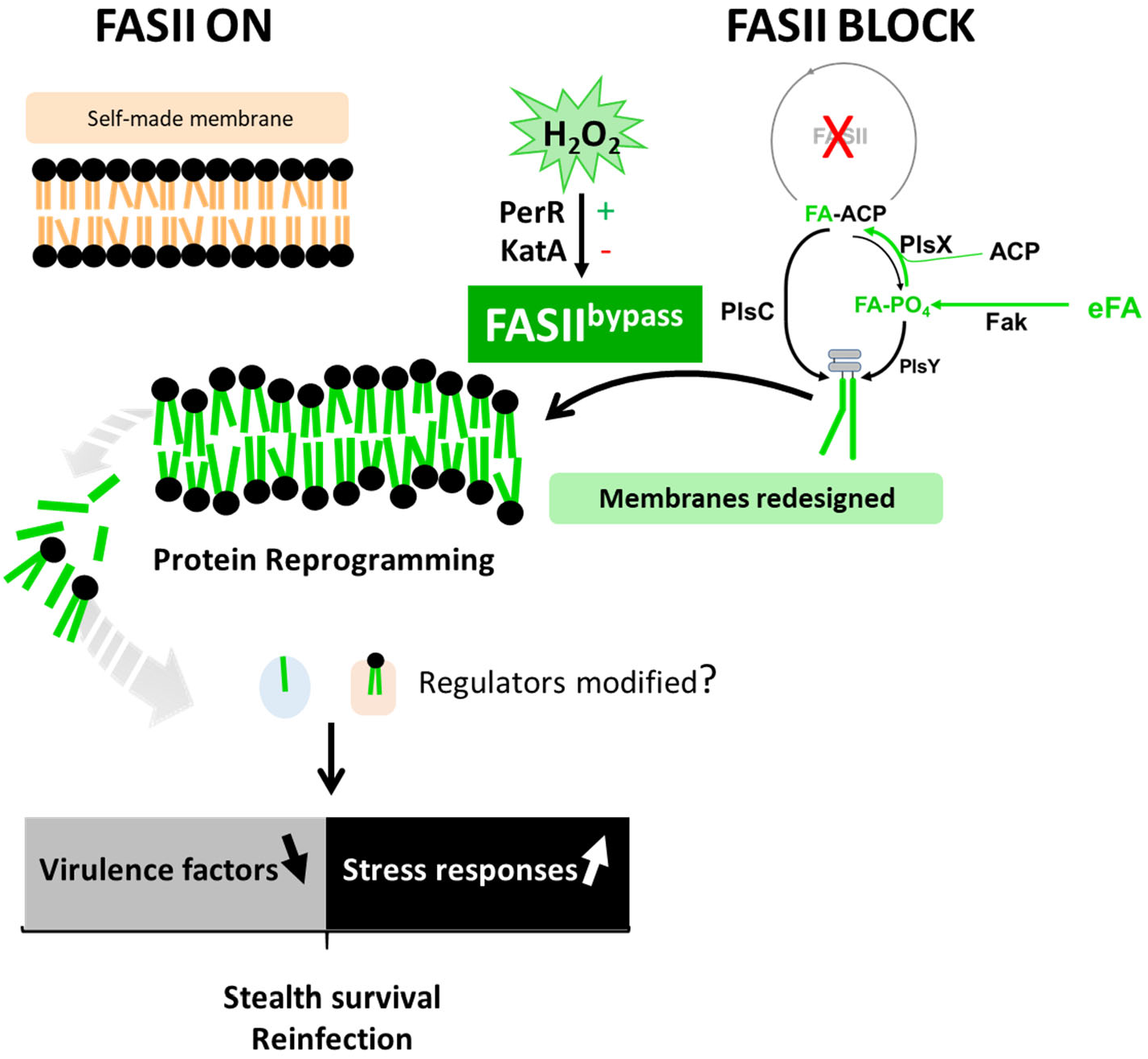
Summary model of anti-FASII adaptation and *S. aureus* fitness. *S. aureus* synthesizes fatty acids (FAs) to produce membrane phospholipids (upper left, FAs in orange). Anti-FASII treatment in SerFA medium promotes FASII bypass, during which exogenous FAs (eFAs, in green) are incorporated and constitute the membrane phospholipid FAs (Kenanian et al., 2019). FASII bypass is accelerated by H_2_O_2_ priming, which requires PerR, but is lower if KatA is present. Anti-FASII adaptation is accompanied by massive changes in protein expression. Membrane perturbation in the new phospholipid environment is proposed to signal protein reprogramming; as reported, membrane FAs or phospholipids may shed internally and bind to regulatory proteins to modulate their function (Schurig-Briccio et al., 2020, Yeo et al., 2023, Lowden et al., 2010, Huang et al., 2022). Decreased virulence factor production may help bacteria escape host immune surveillance (Tuchscherr et al., 2020). Up-regulation of stress response protein levels confers greater ROS tolerance, and could facilitate survival during infection (DeCoursey, 2004). Peroxide priming accelerates FA incorporation and anti-FASII adaptation by a novel process. Anti-FASII treatment would favor emergence of *S. aureus* populations that are at least transiently less infectious, but that may persist in the host.

### Regulator changes during anti-FASII treatment

Various regulators appear to be implicated in anti-FASII adaptation, *e.g*., XdrA or CshA, and to lesser extents HrcA or CcpE. However, no single regulator was found to have a “master” role. In addition to alterations in protein abundance and phosphorylation levels as assessed here, we hypothesize that membrane perturbations, as caused by anti-FASII, alter the intracellular environment, to liberate signaling lipids that modulate regulator activities (Fig. 6). For example, FAs and cardiolipin reportedly affect activities of the *S. aureus* two-component system SaeRS (Schurig-Briccio et al., 2020, Yeo et al., 2023). Virulence factor activity may be blocked by FA binding as shown in *Vibrio cholerae* (Lowden et al., 2010). Interaction of regulators with lipid species released during anti-FASII treatment may be a rapid means to adjust regulator functions for FASII bypass.

### Link between FASII-antibiotic adaptation and *S. aureus* fitness against peroxide stress

We showed that ROS priming stimulates FASII bypass by increasing eFA incorporation (Fig. 4). To our knowledge, this is the first report establishing a connection between oxidative stress and stimulation of FASII antibiotic adaptation, and defines a novel paradigm. This mechanism contrasts to that described in *E. coli*, where ROS priming either lowered proton-motive-force-mediated antibiotic entry and/or induced efflux pump expression, *via* a Suf-controlled Fe-S repair system (Gerstel et al., 2020). In contrast, the H_2_O_2_-stimulated anti-FASII adaptation is mediated by greater, rather than less, eFA incorporation and precocious adaptation, ruling out a role for efflux. A *suf* mutant remained sensitive to peroxide activation, further distinguishing the two mechanisms.

We hypothesize that H_2_O_2_ has a direct role in stimulating FASII bypass: H_2_O_2_–mediated oxidation of FASII initiation enzymes could diminish competition with the FASII bypass system (Fig. 6). We show here that a *katA* mutant, which presumably accumulates H_2_O_2_, shortens adaptation time, while a *perR* mutant, which activates KatA to degrade H_2_O_2_, lengthens adaptation time. Direct or indirect targets of H_2_O_2_ could be involved in disabling FASII to favor FASII bypass; this hypothesis is currently under study.

Host macrophages and neutrophils generate peroxides presumably as a means to control infection (Cole et al., 2014, Mayer-Scholl et al., 2004). *S. aureus* adaptation to anti-FASII is potentiated in these same conditions, which needs to be considered for future use of FASII antibiotics.

### Potential advantages of a fitness reset in FASII-antibiotic-adapted *S. aureus*

Lower virulence, greater stress response, and exclusive eFA utilization may be advantageous to *S. aureus* lifestyle. We note that streptococci and enterococci use FASII bypass even without antibiotics, favoring eFA incorporation during infection (Brinster et al., 2009, Lambert et al., 2023). Although these species employ distinct FASII regulation system from that of *S. aureus* (feedback *versus* feedforward in *S. aureus* (Lambert et al., 2022, Albanesi et al., 2013)) and differ in their *in vivo* lifestyles, they have common infection biotopes, raising the possibility that FASII bypass is advantageous in the course of infection.

Parallels also exist between anti-FASII-adapted *S. aureus*, and small colony variants (SCVs): both emerge more efficiently in oxidative stress (Peyrusson et al., 2022, Painter et al., 2015), and both produce less virulence factors (Tuchscherr et al., 2020, Sendi and Proctor, 2009), which might facilitate bacterial escape from host immune surveillance. In both cases, uneradicated reservoirs of these bacteria might be sources of chronic infection (Fig. 6).

### Potential applications of FASII antibiotics

Although anti-FASII do not eliminate *S. aureus* when compensatory eFAs are available, the present study demonstrates that it reduces *S. aureus* virulence factor production. Anti-FASII synergy with other treatments that prevent adaptation may potentiate its efficacy (Pathania et al., 2021). Here, anti-oxidants delayed anti-FASII adaptation, which may offer perspectives for a bi-therapy approach to eliminate *S. aureus* with reduced virulence.

## Materials and Methods

### Strains, media, and growth conditions

Experiments were performed using *S. aureus* SAUSA300_FPR3757 JE2 strain, referred to as USA300, and transposon insertion derivative strains from the Nebraska mutant library ((Fey et al., 2013); generously supplied by BEI Resources NIAID, NIH, USA). All strains were confirmed for transposon insertion by PCR (see **Table S4** for strains and primers used). Cultures were grown aerobically at 37°C. Solid and liquid growth media were based on BHI as follows: no additives (BHI), containing 0.5 mM FAs (BHI-FA, with an equimolar mixture of 0.17 mM each C14, C16, C18:1 [Larodan, Sweden]), and BHI-FA containing 10% newborn calf serum (Ser-FA; Eurobio Scientific, France [Fr]). Where specified, the anti-FASII triclosan (McMurry et al., 1998) was added at 0.25 µg/ml in media without serum, and at 0.5 µg/ml in media containing serum, to respectively give BHI-Tric, FA-Tric, or SerFA-Tric, as described (Morvan et al., 2017, Kenanian et al., 2019, Morvan et al., 2016). H_2_O_2_ (0.5 mM final concentration) was added to SerFA precultures when indicated. The anti-FASII AFN-1252 (Karlowsky et al., 2009) was used in SerFA at 0.5 µg/ml (SerFA-AFN) as described (Kenanian et al., 2019). For most experiments, *S. aureus* USA300 was streaked on solid BHI medium, and independent colonies were used to inoculate overnight BHI pre-cultures. For proteomic and phosphoproteomic studies, cultures were inoculated at a starting OD_600_ = 0.1.

### Nanopore sequencing

Six single colonies of USA00 were resuspended in SerFA and grown overnight. Cultures were then diluted in SerFA, SerFA-AFN-1252, 10% mouse serum, or 10% mouse serum containing AFN-1252, and grown to OD_600_ = 1. Whole chromosomal DNA was prepared from the independent cultures as described, and outsourced for nanopore sequencing (Eurofins, Germany). Whole genome DNA sequences were presented as circularized genomes, and entered in the Mendeley database doi:10.17632/grt4htck9k.1. Sequences were compared by performing full genome alignments https://dgenies.toulouse.inra.fr/ 09/02/2023.

### Proteomics preparation

Adaptation to anti-FASII varies with growth media: BHI-Tric-grown and FA-Tric-grown *S. aureus* do not adapt in the time periods tested, (high frequency adaptive mutations arise with a delay in FA-Tric; (Morvan et al., 2016)), whereas SerFA-Tric-grown *S. aureus* adapt without mutation after an initial latency period (6-8 hours, depending on growth conditions) (Kenanian et al., 2019). Kinetics experiments were performed on USA300 to determine the protein changes associated with FA-Tric and SerFA-Tric anti-FASII adaptation. Cultures for each condition were prepared as independent quadruplicates. For each sample, BHI precultures were diluted and shifted to the specified medium starting cultures at OD_600_ = 0.1. Control cultures in BHI, BHI-FA and BHI-SerFA were grown to OD_600_ = ∼1. BHI-Tric cultures (no added FAs) were collected at 6 h. USA300 samples grown in FA-Tric and SerFA-Tric were collected at 2, 4, 6, 8 and 10 h post-antibiotic addition (see Fig. 1A and **Table S1** for growth conditions and complete data). For each sample, 20 OD_600_ units culture equivalent was collected and centrifuged for 10 min at 4°C at 8000 rpm. Pellets were washed twice in Tris 10mM pH7.0 containing 0.02% TritonX-100, and Halt^TM^ Protease & Phosphatase Inhibitor Cocktail (100X) (Thermo Scientific, Fr). Pellets were then resuspended in 650 µl washing buffer, mixed with 0.1 mm silica beads and subjected to 3 cycles of vigorous shaking (Fast-Prep-24, MP-Bio, Fr). After 10 min centrifugation at 10000 rpm, supernatants were recovered and stored at −80°C prior to analyses.

Protein extractions, LC-MS/MS analyses, and bioinformatics, and statistical data analyses were done as described in detail (Bednarz et al., 2021). The reference genome GenBank Nucleotide accession code NC_007793.1 was used for protein annotation. The bioinformatic tools used for proteomics analysis (X!TandemPipeline C++, MassChroQ, MCQR) are open and free resources available in the following repository: https://forgemia.inrae.fr/pappso. The mass spectrometry proteomics data was deposited to the ProteomeXchange Consortium (http://proteomecentral.proteomexchange.org) via the PRIDE partner repository with the dataset identifier PPXD034256.

### Proteome data analyses

Protein abundance differences were detected by ANOVA tests for all methods used (spectral counting, SC; extracted ion chromatograms, XIC; and peak counting, PC). The abundance of a protein was considered significantly variable when the adjusted p-value was <0.05. Proteins showing significant statistical abundance differences were represented on heat maps normalized by using GraphPad Prism 9.5.1 (Bednarz et al., 2021). K-means clustering analysis was performed using RStudio. Proteins whose abundancies were significantly different in one or more conditions were manually curated and classified according to functional groups.

### Phosphoproteome preparation

USA300 cultures were prepared in BHI and SerFA and harvested at OD_600_ = ∼1. SerFA-Tric cultures were harvested at 6h and 10 h post-treatment. Samples were prepared independently from those in the proteomics study, as independent biological triplicates, and treated as in proteomics studies, except that we collected the equivalent of 50 OD_600_ units. Bacteria were processed as for proteome extraction except that Tris was replaced by triethylammonium bicarbonate (50 mM; *n.b.* Tris interferes with dimethyl tag labelling) containing antiprotease and antiphosphatase at the recommended concentrations. Lysed bacteria after Fast-Prep were centrifuged 15 min at 12000 rpm supernatants containing soluble proteins were kept at −80°C before use. Protein concentrations were determined by the Bradford method.

One mg protein samples were evaporated and resuspended in 1 ml 5 % formic acid and dimethyl-tag labeled as described (Boersema et al., 2009). Briefly, differential on-column labeling of peptide amine groups (NH_2_) created dimethyl labels leading to mass shifts of +28.0313 Da for the peptides from SerFA 3 h samples, +32.0564 Da for the peptides from SerFA-Tric 6 h samples and +36.0757 for the peptides from SerFA-Tric 10 h samples respectively. The three samples were mixed and then submitted to six rounds of phosphopeptide enrichment with 5mg TiO_2_ beads/mg of protein (Titansphere Phos-TiO, GL Sciences Inc., Netherlands) as described (Soufi et al., 2018).

LC-MS/MS analyses of samples were done using an Ultimate 3000 nano-RSLC coupled on line with a Q Exactive HF mass spectrometer (Thermo Scientific, San Jose California). 1 µL of each sample was loaded on a C18 Acclaim PepMap100 trap-column 300 µm inner diameter (ID) x 5 mm, 5 µm, 100Å, (Thermo Scientific) for 3.0 minutes at 20 µL/min with 2% acetonitrile (ACN), 0.05% TFA in H_2_O and then separated on a C18 Acclaim Pepmap100 nano-column, 50 cm x 75 µm ID, 2 µm, 100 Å (Thermo Scientific) with a 100 minute linear gradient from 3.2% to 20% buffer B (A: 0.1% FA in H_2_O, B: 0.1% FA in ACN), from 20 to 32% of B in 20 min and then from 32 to 90% of B in 2 min, hold for 10 min and returned to the initial conditions. The flow rate was 300 nL/min.

Labeled peptides were analyzed with top15 higher energy collisional dissociation (HCD) method: MS data were acquired in a data dependent strategy selecting the fragmentation events based on the 15 most abundant precursor ions in the survey scan (m/z range from 350 to 1650). The resolution of the survey scan was 120,000 at m/z 200 Th and for MS/MS scan the resolution was set to 15,000 at m/z 200 Th. For HCD acquisition, the collision energy = 27 and the isolation width is of 1.4 m/z. The precursors with unknown charge state, charge state of 1 and 5 or greater than 5 were excluded. Peptides selected for MS/MS acquisition were then placed on an exclusion list for 20 s using the dynamic exclusion mode to limit duplicate spectra. The mass spectrometry proteomics data have been deposited to the Center for Computational Mass Spectrometry repository (University of California, San Diego) *via* the MassIVE tool with the dataset identifier MassIVE MSV000089781 (accessible at http://massive.ucsd.edu/ProteoSAFe/status.jsp?task=db3f5002e32e4a47a71996358bc6ae8c).

### Phosphoproteome data analyses

Proteins were identified by database searching using SequestHT with Proteome Discoverer 2.5 software (Thermo Scientific, Fr) against the Uniprot *S. aureus* USA300 database (2020-01 release, 2607 sequences). Precursor mass tolerance was set at 10 ppm and fragment mass tolerance was set at 0.02 Da, and up to 2 missed cleavages were allowed. Oxidation (M), acetylation (Protein N-terminus), and phosphorylation (S, T, Y) were set as variable modifications. The differentially dimethyl-labeled peptides in primary amino groups K and N-ter (see above), and carbamidomethylation (C) were set as fixed modifications. Peptides and proteins were filtered with a false discovery rate (FDR) at 1% using the Percolator tool (The et al., 2016). Protein quantitation was performed with precursor ions quantifier node in Proteome Discoverer 2.5 software, peptide and protein quantitation based on pairwise ratios and t-test statistical validation.

### Macrophage adhesion

Confluent cell lawns of THP-1 macrophages were prepared as described to obtain confluent cell lawns of 3×10^5^ cells per well (Bourrel et al., 2022). *S. aureus* was pre-cultured in SerFA medium and then subcultured overnight in SerFA medium without or with AFN-1252. The next day, bacteria were subcultured in the same corresponding fresh media to OD_600_ =1-2. Bacteria were then washed twice with PBS buffer, and diluted in RPMI GluMax (Gibco, France) to obtain an MOI = 1. One ml of the dilution was added in each well in 24-well plates. Plates were centrifuged at 1000 rpm for 5 minutes to sediment bacteria, and then incubated at 4°C for 1h. After incubation, wells were washed 3 times with PBS, then 1ml of sterile cold water was added to each well and left for 5min at room temperature. Finally, the cells and bacteria were scraped from the surface and CFU were determined on SerFA agar plates. Controls without macrophage were done in parallel. Results are derived from 5 biological replicates, and are presented as the mean ± standard error of the mean (SEM) using GraphPad Prism 9.5.1 (San Diego, Ca). CFU counts were compared by paired t-test (P<0.05).

### Exoprotein activity assays

SerFA day precultures were diluted into SerFA without or with either triclosan (SerFA-Tric) and AFN-1252 (SerFA-AFN). Solid medium as prepared for exoprotein detection were then spotted with resulting overnight saturated cultures or culture supernatants as indicated. For nuclease detection, DNase agar (Oxoid, Thermo Scientific, Fr) containing toluidine blue O 0.05 g/L (Sigma, Fr) was prepared as described (Zierdt and Golde, 1970). Culture supernatants from test samples were heated to 80°C for 10 m, and then spotted (10 µl) on plates. Photographs were taken after overnight incubation at 37°C. Contrast was uniformly enhanced by Photoshop to visualize pink halos indicating nuclease activity. To measure protease activity, powdered skim milk 50 g/L was added to autoclaved 1% non-nutrient agar (Invitrogen) as described (Park et al., 2012). Cultures were spotted (10 µl) on plates and allowed to dry, then incubated at 37°C for 72 h and photographed. Lipase activity was assayed in medium comprising 1% non-nutrient agar (Invitrogen) to which was added 2.5% olive oil and 0.001% Rhodamine B (starting from a Rhodamine B stock solution of 1 mg per ml in water; Sigma-Aldrich, Fr) as described (Araiza-Villanueva et al., 2019). To improve olive oil emulsification, NaCl was added to medium (1M final concentration), followed by vigorous shaking just prior to plate preparation. Supernatants from overnight cultures were sterile-filtered through a 0.2 µm membrane syringe filter (Pall Corporation, Michigan). Supernatants (10 µl) were deposited in holes pierced in solid medium. After 24 h incubation at 37°C plates were visualized under UV light at 312 nm and photographed. Hemolytic activity was assayed on *S. aureus* sterile-filtered supernatants that were prepared from overnight cultures. Aliquots (10 µl) were spotted on 5% sheep blood agar plates (BioMérieux SA, Fr). After drying, plates were incubated at 37°C overnight, and photographed.

### H_2_O_2_ resistance of non-treated and anti-FASII-adapted *S. aureus*

*S. aureus* non-treated or AFN-adapted cells (see above) were diluted to OD_600_= 0.005 in SerFA and grown for 1 h at 37°C, after which H_2_O_2_ (0.5mM final concentration) was added or not to cultures, and allowed to grow for 5 h. Growth kinetics was followed by OD_600_ determinations, and CFUs were determined at the final time point. Plates were photographed after overnight incubation at 37°C.

### H_2_O_2_ and PMS pre-treatment

For pre-adaptation in hydrogen peroxide or in PMS, *S. aureus* were precultured in SerFA and then subcultured in SerFA containing or not 0.5 mM H_2_O_2_ (final concentration) or PMS (20-50 µM) for 16 h at 37°C. Cultures were diluted to OD_600_ = 0.1 in SerFA or SerFA medium containing AFN-1252 (SerFA-AFN), and growth was monitored.

### *G. mellonella* infection by anti-FASII-adapted and untreated *S. aureus*

Killing capacities and CFUs of anti-FASII-adapted and untreated *S. aureus* USA300 were compared in the *G. mellonella* insect model (Sheehan et al., 2019). *G. mellonella* larvae were reared on beeswax and pollen in sealed containers with a wire mesh lid permitting aeration. The rearing container was stored at 27°C temperature in a humidified incubator in our laboratory facilities. Fifth instar larvae weighing ∼250 mg were subjected to starvation for 24 h at 27°C and 1-2 h at 37°C prior to infection. *S. aureus* SerFA cultures were subcultured in SerFA, and SerFA plus AFN-1252 (0.5 µg/ml; SerFA-AFN) for overnight growth. Resulting cultures were then diluted to 0.1 and grown in respective media to OD_600_ = 1. Based on previous CFU determinations, each culture was pelleted, washed once in PBS, and then resuspended in PBS to obtain 10^8^ CFU/ml. Ten µl (10^6^ CFUs) was administered by injection in the fourth proleg. To follow larvae mortality following *S. aureus* infection, 20 larvae were inoculated using 3 independent cultures per culture condition (totaling 60 insects injected with SerFA, and 60 with SerFA-AFN), and 20 larvae were inoculated with PBS buffer for the control. Surviving larvae were counted every 24 h for 72 hours. To monitor *S. aureus* bacterial counts post-infection, the experiment above was repeated, and surviving larvae (9 total per condition, treated in 3 groups from independent biological replicates) were sacrificed until there were no surviving larvae left. CFUs were also determined from newly-dead larvae. To determine *S. aureus* CFUs, surviving larvae were chilled in ice for 30 min, and then crushed in 1 ml sterile deionized water. The resulting content was vortexed for ∼30 sec. Five µl of 10-fold dilutions were prepared in sterile water and spotted on SerFA and SerFA-AFN solid media. CFUs were determined after overnight incubation at 37°C. The detection threshold was 10^3^ CFU per insect. Results of the 3 biologically independent experiments were pooled for presentation of insect mortality, and similarly for CFUs as done in the second independent experiment. Results are presented as the mean ± standard deviation (SD) using GraphPad Prism 9.5.1 (San Diego, Ca). CFU counts were compared by nonparametric t-test (Mann-Whitney U) (P<0.05). Note that *G. mellonella* naturally harbor enterococci (Allonsius et al., 2019). In contrast to *S. aureus*, enterococci are catalase-negative on routine plate tests (*n.b.*, some enterococci produce catalase upon heme addition). *S. aureus* were thus differentiated from enterococci on plates used for CFU enumeration by their catalase-positive response (*i.e.*, bubble formation) upon application of a 3% hydrogen peroxide solution. Enterococci did not produce bubbles.

## Acknowledgments

The Nebraska Transposon Mutant Library strains were generously provided by BEI Resources, NIAID, NIH, USA, and by Dr. P. Fey (Nebraska Medical Center, Omaha, USA). We acknowledge Micalis colleagues V. Sanchis and C. Nielson-LeRoux for valuable advice on the *G. mellonella* infection model. We gratefully acknowledge our colleagues for their skillful technical assistance: C. Buisson (Micalis) for insectarium and insect model, L. Dupont (Micalis) for proteomic sample preparations, J. Guignot (Institut Cochin) for adhesion experiments, and F. Delolme (Université de Lyon) for phosphoproteome studies. We thank P. Gaudu (Micalis) for valuable discussions.

## Data Availability

All relevant data are within the manuscript, supporting information files, and depositories.

