## Supplementary Figures 1-5 for "Oxidative Stress is Intrinsic to Staphylococcal Adaptation to Fatty Acid Synthesis Antibiotics"

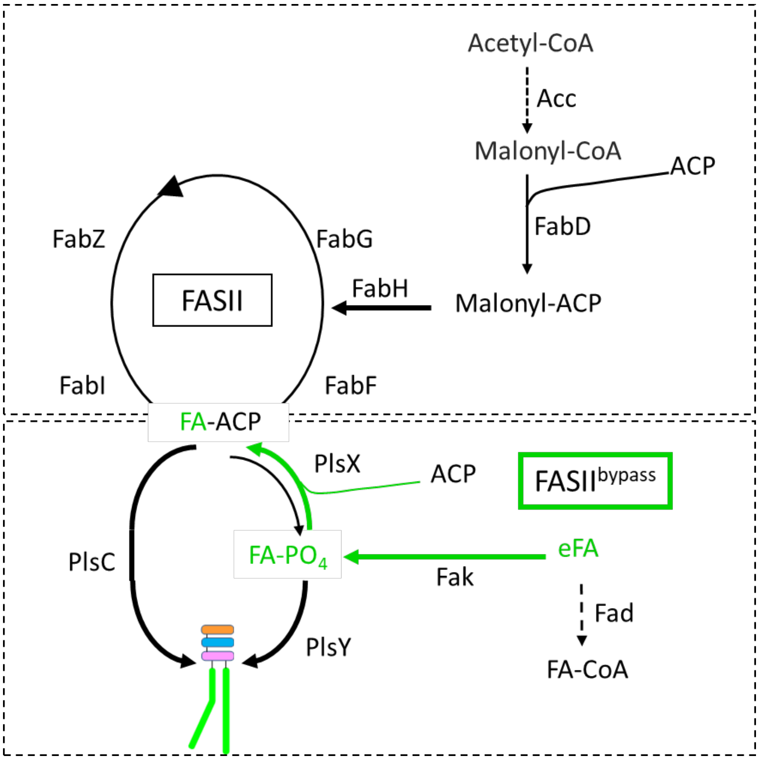


**Fig S1. Schematic model of FASII and FASII bypass in *S. aureus*. Upper:** *S. aureus* synthesizes fatty acids (FAs) *via* the fatty acid synthesis pathway FASII (boxed in black), comprising 3 dedicated initiation enzymes (upper right), and 4 enzymes that constitute the recursive FASII cycle (for review, (Zhang and Rock, 2008))**.** The final product, FA-ACP (ACP, acyl carrier protein), provides FAs for transfer to a glycerol phosphate backbone for phospholipid synthesis. **Lower:** FASII bypass (boxed in green) can take over when FASII is inhibited. Environmental FAs are phosphorylated by fatty acid kinase (Fak; (Parsons et al., 2014)). Both FASII and FASII bypass products lead to phospholipid synthesis: Starting from FA-ACP (the FASII product), PlsX and PlsY catalyze FA attachment to position 1 of the glycerolphosphate backbone. PlsC then joins a second FA to position 2 of the mono. Starting from eFAs, the resulting phosphorylated FA (FA-PO$_4$) is used by PlsY to join a FA to position 1. Reverse activity of PlsX generates FA-ACP, which is the substrate for PlsC to join the FA to position 2 on the lysophosphatidic acid. eFAs are also substrates for Fad (fatty acid degradation) enzymes, dotted arrow, which are encoded by *S. aureus*, although their activity is not yet demonstrated.

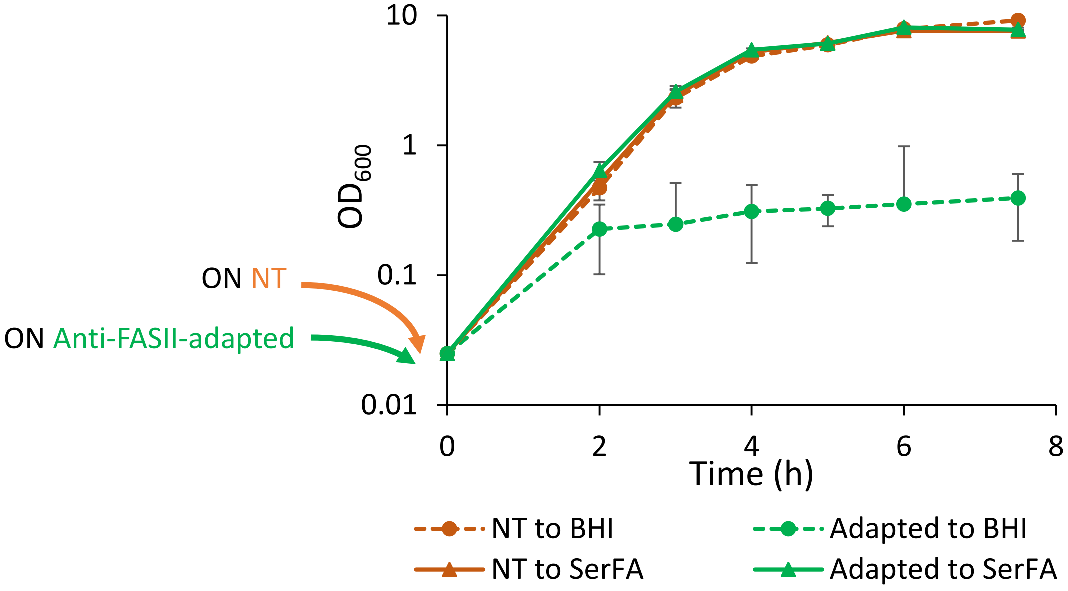


**Fig. S2. Regrowth of non-treated and anti-FASII-adapted *S. aureus* USA300 cultures varies according to medium.** Overnight (ON) non-treated (NT) and anti-FASII adapted *S. aureus* USA300 cultures were prepared in SerFA containing or not anti-FASII AFN-1252. They were then diluted to $OD_{600}$ 0.025 in SerFA or BHI medium without selection. Growth of cultures was monitored. Samples are as indicated above. Shown are the mean and standard deviation of 3 biological replicates.

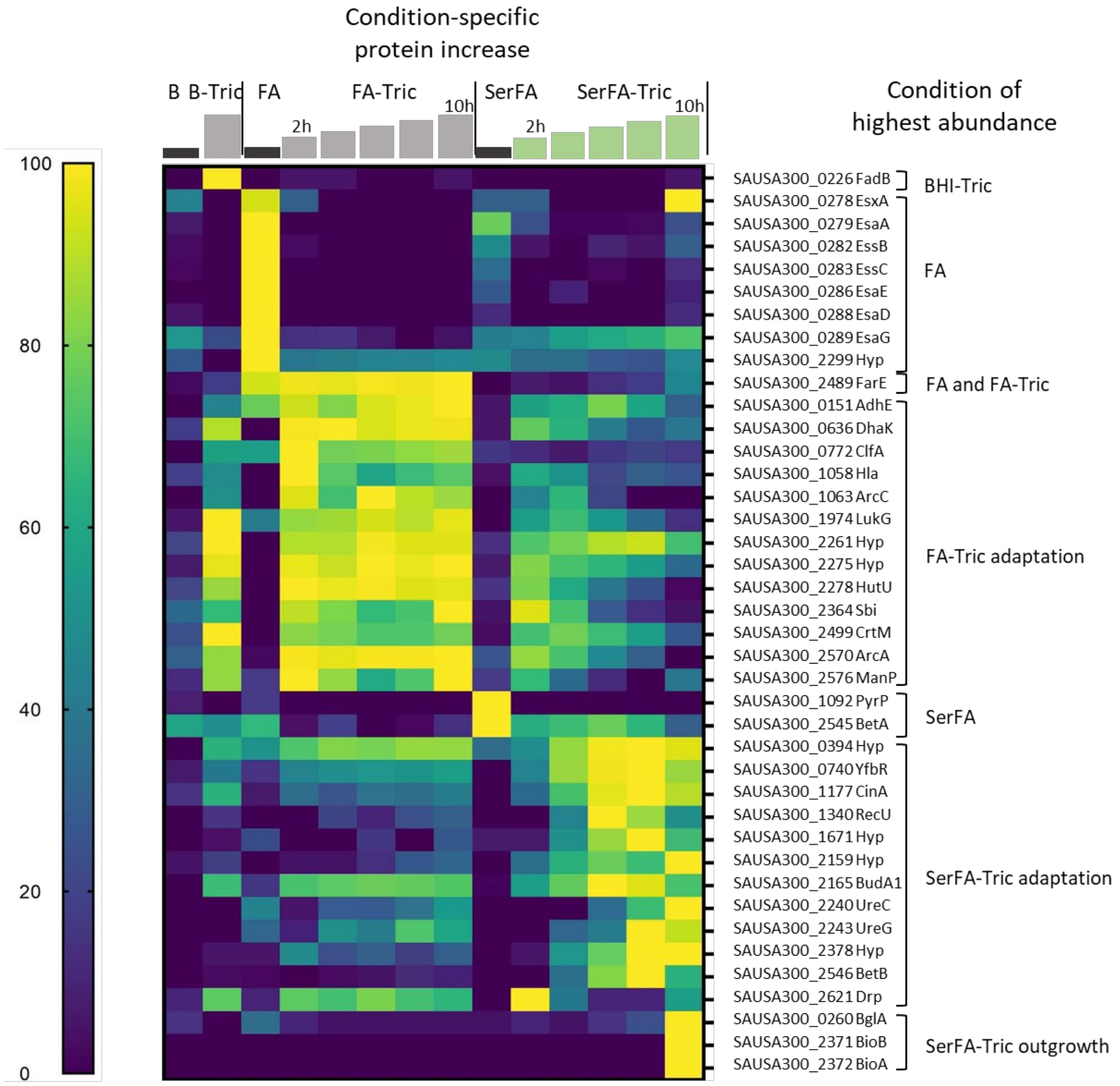


**Fig. S3. Heat map of condition-specific highly expressed proteins.** Gene loci and encoded proteins at right show the highest differential expression in conditions indicated after each bracket. FA-Tric and SerFA-Tric kinetics samples were each analyzed as a group regardless of the time of increased expression. Hyp, hypothetical protein. The heat map scale (at left) is determined relative to weighted value for each protein (navy, down-represented; yellow, up-represented). Sampling times (h) above steps correspond to 2, 4, 6, 8, and 10 h for FA-Tric and SerFA-Tric. Green steps corresponds to the adaptation condition.

### FASII and Phospholipid enzymes

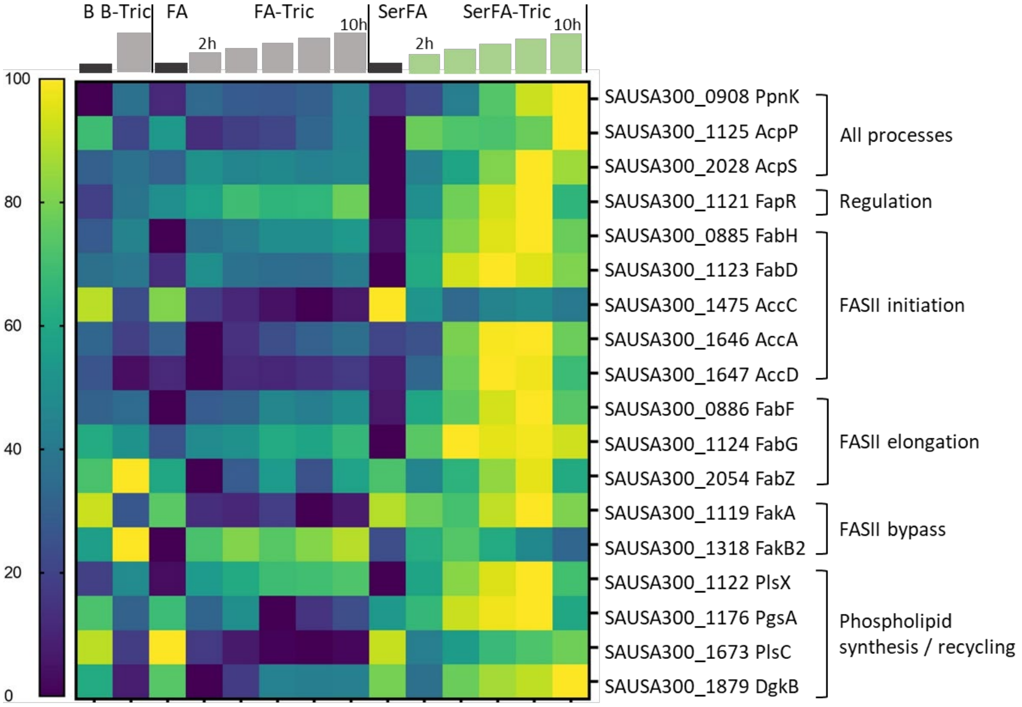


**Fig. S4. Heat map of FASII and phospholipid synthesis and recycling functions whose expression is affected in SerFA-Tric growth.** Proteins were compiled based on confirmed FASII and phospholipid synthesis and recycling proteins. Proteins unaffected or nondetected in any condition are not included. Results are limited to SerFA (control) and SerFA-Tric adaptation conditions. Gene names and functional categories are at right. Incremental time points for SerFA-Tric 2, 4, 6, 8, and 10 h samples are represented by steps. Correspondence between color and expression is determined relative to weighted value for each protein, as on scale at left (navy, down-represented; yellow, up-represented).

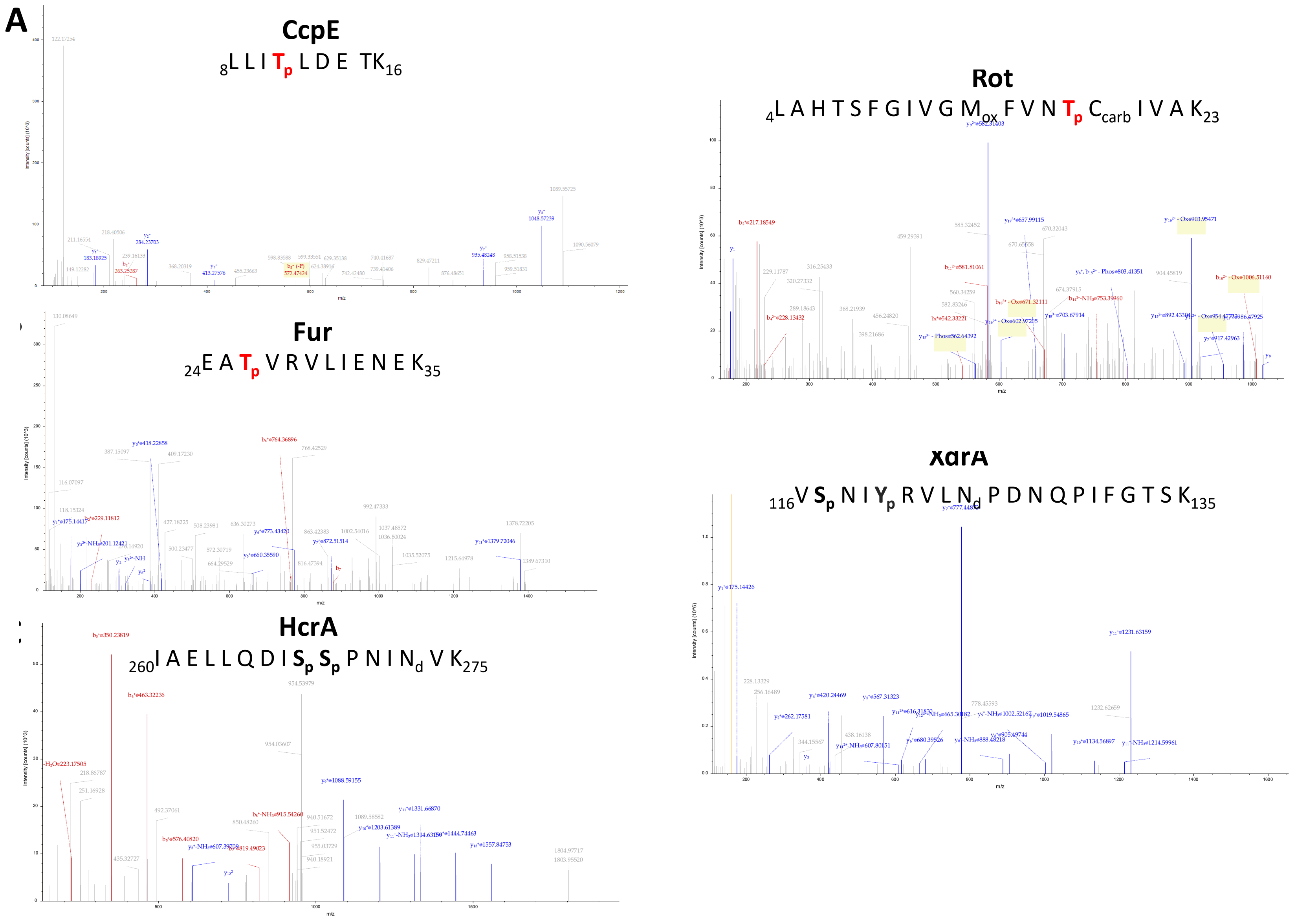


**Fig. S5. Tandem Mass spectra of identifying differentially expressed phosphorylation sites of CcpE (A), Fur (B), HcrA (C), Rot (D) and XdrA (E) proteins.** The identified phosphorylation sites are indicated in bold red when localization is validated and in bold black when 2 phosphorylation sites are possible. d: deamidation, carb: carbamidomethylation, ox: oxidation, p: phosphorylation. All peptides are labeled by dimethyl tag on peptide N-term and K.
